# Comparative Analysis, Diversification and Functional Validation of Plant Nucleotide-Binding Site Domain Genes

**DOI:** 10.1101/2023.08.09.552572

**Authors:** Athar Hussain, Aqsa Anwer Khan, Muhammad Qasim Aslam, Aquib Nazar, Nadir Zaman, Ayesha Amin, Muhammad Arslan Mahmood, M. Shahid Mukhtar, Hafiz Ubaid Ur Rahman, Muhammed Farooq, Muhammed Saeed, Imran Amin, Shahid Mansoor

## Abstract

Nucleotide-binding site (NBS) domain genes are one of a superfamily of resistance genes involved in plant responses to pathogens. The current study identified presumably identified 12,820 NBS-containing genes across 34 species covering from mosses to monocots and dicots. These identified genes classified into 168 classes with several novel domain architectures patterns encompassing significant diversity among plant species. Several classical (NBS, NBS-LRR, TIR-NBS, TIR-NBS-LRR etc.) and species-specific structural patterns (TIR-NBS-TIR-Cupin_1-Cupin_1, TIR-NBS-Prenyltransf, Sugar_tr-NBS etc.) were discovered. We observed 603 orthogroups (OGs) with some core (most common orthogroups; OG_0_, OG_1_, OG_2_ etc.) and unique (highly specific to species; OG_80_, OG_82_ etc) OGs with tandem duplications. The expression profiling presented the putative upregulation of OG_2_, OG_6,_ and OG_15_ in different tissues under various biotic and abiotic stresses in susceptible and tolerant plants to CLCuD. The genetic variation between susceptible (Coker 312) and tolerant (Mac7) *G. hirsutum* accessions identified several unique variants in NBS genes of Mac7 (6,583 varaints) and Coker312 (5,173 variants). The protein-ligand and proteins-protein interaction showed a strong interaction of some putative NBS proteins with ADP/ATP and different core proteins of cotton leaf curl disease virus. The silencing of *GaNBS* (OG_2_) in resistant cotton through virus-induced gene silencing (VIGS) demonstrated its putative role in virus tittering. The presented study will be further helpful to understand the plant adaptation mechanism.

## Introduction

Gene duplication and loss events constitute the main factors of gene family evolution [1]. Duplications occur through two major processes, whole-genome duplication (WGD) and small-scale duplications (SSD), including tandem, segmental, and transposon-mediated duplications [2]. They appear to be distinct modes of expansion, since gene families evolving through WGDs rarely experience SSD events [3] and maintain gene family expansion.

The R gene-mediated defense responses are one of the major sources of resistance against plant pathogens including viruses [4] and nucleotide-binding leucine-rich repeat (NLR) is one of the major families of R gene. Similar to animal NLRs, plant NLRs are modular proteins that generally consist of three building blocks: an N-terminal domain, the central NB-ARC domain (named after Nucleotide-Binding adaptor shared with APAF-1, plant resistance proteins, and CED-4), and a C-terminal LRR (leucine-rich repeats) domain [5, 6]. The central domain of animal NLRs is also known as the NACHT domain [7] which is structurally similar to the plant NB-ARC domain but distinctive of animal NLRs [8, 9]. The utilization of either a TOLL/interleukin 1 receptor (TIR) domain or a coiled-coil (CC) domain at the N-terminus is a plant-NLR-specific feature and defines two major types of plant NLRs termed the TIR-type NLRs (TNLs) and the CC-type NLRs (CNLs), respectively [10, 11]. However, it is often challenging to specify structures of N-terminal domains for a significant proportion of plant NLRs due to their structural diversity and lack of significant homology to validated protein structures [12]. Thus, NLRs containing an N-terminus other than the TIR domain are sometimes designated as non-TIR-type NLRs (nTNLs) as a distinction to TNLs.

The NLR family has massively expanded in many plants. The massive expansions render the NLR family one of the largest and most variable plant protein families [13]. This contrasts with the vertebrate NLR repertoires, typically comprising ca. 20 members [14, 15]. Detailed genome-wide surveys, database mining, and degenerate PCR approaches for the species whose genome sequences are currently not available to contribute to refining an overview of the NLR repertoires in various plant species [16]. Most of the plant genomes surveyed so far have a large NLR repertoire with up to 459 genes in wine grapes [17]. Interestingly, the bryophyte *Physcomitrella patens* and the lycophyte *Selaginella moellendorffii* which represent the ancestral land plant lineages seem to have a relatively small NLR repertoire of ∼25 and ∼2 NLRs respectively, suggesting that the gene expansion has occurred mainly in flowering plants. It was recently shown that numerous microRNAs target nucleotide sequences encoding conserved motifs of NLRs (e.g., P-loop) in many flowering plants [18]. Thus it is hypothesized that such a bulk control of NLR transcripts may allow a plant species to maintain large NLR repertoires without depletion of functional NLR loci [19, 20] since microRNA-mediated transcriptional suppression of NLR transcripts could compensate for the fitness costs related to maintenance of NLRs [18, 21].

Currently, in Pakistan, almost all cotton varieties of *Gossypium. hirsutum* are vulnerable to cotton leaf curl disease (CLCuD) [22]. CLCuD is caused by Begomoviruses (family: Geminiviridae) and transmitted by insect vector, whitefly (*Bemisia tabaci*).. *G. arboreum,* also known as “Desi cotton,” represents a high level of resistance against insect pests and diseases including CLCuD, while Mac7 (*G. hirsutum*) is highly tolerant, and Coker-312 (*G. hirsutum*) is highly susceptible [23, 24]. NBS-LRR is the main classes of resistance genes that showed response to viruses [25, 26]. Thus, this study aimed to evaluate the NBS domain associated with host plant resistance genes, a potential source of disease resistance.

## Methodology

### Genome assemblies and data collection

In the current study, we have selected 39 land plants ranging from green algae to higher plants families including Amborellaceae, Brassicaceae, Poaceae, Citrus, Cucurbitaceae, Malvaceae, Marchaceae, Fabaceae, Nelumbonaceace, Salicaceae, Rosaceae, and Araceae families. In addition, the selection of plants was also made based on ploidy level (haploid, diploid, and tetraploid) for further detailed evolutionary study. The latest genome assemblies (Table S1). were downloaded from publicly available respective genome databases, NCBI, Phytozome and Plaza genome databases[27, 28].

### Identification, classification, and comparison among land plants

In order to screen the NBS (NB-NRC) domain-containing genes, the *PfamScan.pl* HMM search script was used with e-value (1.1e-50) using background Pfam-A_hmm model [29]. All genes having NB-ARC domains were considered NBS genes and filtered. In addition, the additional associated decoy domains were also observed through the domain architecture of NBS genes by following the Hussain et al. [30] classification method. In this classification system, similar domain-architecture-bearing genes were placed under the same classes. Furthermore, comprehensive comparison of classes was also made among land plants.

### Evolutionary study; Species-based phylogenetic tree, orthogrouping, and duplication analysis

In order to provide a deep understanding of the evolution and diversification of NBS genes in land plants, we used OrthoFinder package tools [31]. In this package, the DIAMOND tool was used for fast sequence similarity searches among NBS sequences. The clustering of genes was done using the MCL clustering algorithm. The orthologs and orthogrouping were carried out with DendroBLAST [32]. For multiple sequence alignment, MAFTT 7.0 was used, and the aligned file was subjected to species-based phylogenetic tree construction using FastME 2.0 version. A gene-based phylogenetic tree was also constructed by the maximum likelihood algorithm in FastTreeMP with a 1000 bootstrap value.

### Transcriptomic analyses of NBS genes

To ascertain the differential expression and responsiveness of NBS genes in various tissues and stresses, we have retrieved RNA-seq data (Arabidopsis, maize, soybean, upland cotton, and wild cotton) from different databases including RNA-seq database Arabidopsis RNA-seq database[33], maize RNA-seq database, cotton RNA-seq database, soybean RNA-seq database, Cotton Functional Genomics Database (CottonFGD)[34] and Cottongen database [35]. Besides the databases, we also collected additional RNA-seq data from NCBI BioProjects (Bio-projects PRJNA490626 and PRJNA594268, (PRJNA390823) and PRJNA398803)[36–38]. The RNA-seq data is categorized into three types tissue-specific (leaf, stem, flower, pollen, endosperm, pollen, and seed etc.), abiotic stress-specific (dehydration, cold, drought, heat, dark, osmotic, salt, wounding etc.) and biotic-stress specific (*Blumeria graminis, Botrytis cereal, Collettrichum tofieldiae, Heterodera schachti nematodes, bacterial strain, Fusarium graminearum, Rhizotonia solani etc.*) expression profiling. The RNA-seq data was processed through transcriptomic pipelines. The final heat map was d’rawn using the TBTool package under heatmap construction with a Log2Base value of FPKM [39].

### Genetic marker prediction in susceptible and tolerant cotton accessions

In order to find important genetic markers in NBS genes, we have selected two contrast accession of *G. hirsutum* cotton. The Mac7, a highly tolerant *G. hirsutum* to CLCuD, and Coker 312, a highly susceptible accession to CLCuD. The whole genome resequencing data of these plants were collected from NCBI BioProject: PRJNA756435 [40] and PRJNA542238 respectively. The NGS Raw read were mapped to NBS genic regions of *G. hirsutum* TM-1 reference genome and identified variants (SNPs/InDels). Furthermore, the identified variants were annotated and characterized based on variant type. The variants associated with genes were further compared between the two-accession using Venny-Bioinfo Tool.

### Gene ontology, KEGG pathways, and *cis*-regulatory elements analysis

To find the functional analysis of NBS genes, we performed gene ontology and KEGG pathway enrichment analysis using the Gene Ontology resource and KEGG PATHWAY Database[41]. The protein features including a number of amino acids, molecular weight, theoretical pI (isoelectric point), and Grand Average Hydropathy (GRAVY), of *G. hirsutum* was studied using ProtParam tool – Expasy tool. For the identification of the cis-regulatory elements of NBS genes in *G. hirsutum*, we have retrieved 2000b upstream of NBS genes and subjected to Plant CARE databases[42].

### Protein modeling, molecular docking, and target gene mining

To determine the NBS protein’s structure and its binding activity to CLCuD viral proteins [43] and with ATP and ADP molecule, we have used the proteins modeling and molecular docking approaches. Based on the biotic stress RNA-seq, we have selected NBS genes (upregulated in tolerant; Mac7, while downregulated in susceptible, Coker 312) from wild cotton (*G. arboreum*; naturally resistant to CLCuD) and upland cotton (*G. hirsutum*, susceptible to CLCuD) data. The 3D structure of Viral and host proteins was predicted using I-TASSER server with default values[44]. The molecular docking was done using Auto-dock Vina[45] and MDock web-based server [46]. The 2D and 3D structures were visualized with LigPlus and discovery studio, respectively [47]. The binding Gibs free energy was calculated with the ClusPro tool [48]. For further functional analysis, we selected Gar06G24920 (OG_2_ member), previously reported as differentially expressed gene under CLCuD in *G. arboreum*.

### VIGS assay and CLCuD inoculation through grafting and viruliferous whitefly exposure

In order to define the function of OG2 genes (*Gar06G24920_OG2*), we have selected a naturally immune *Gossypium sp*, *G. arboreum* variety FDH-228 and used VIGS (virus inducing gene silencing) approaches to silence the selected gene (see supplementary file for detail procedure). The silence plant is then treated with CLCuD through grafting (see supplementary file for detail procedure).

### RNA extraction and qPCR assay

To asses the VIGS and the viral titer, a total RNA was extreacted from VIGs-silenced and control plants. For RNA isolation, leaf tissue samples were taken from the upper three leaves of each plant and RNA was isolated using Trizol (Invitrogen). Purified RNA was used to synthesize cDNA using Revert Aid’s first-strand cDNA synthesis kit (Thermo Scientific, USA). RT-qPCR was performed on at least three independent samples of VIGS and TRV:00 inoculated control plants to measure gene silencing. The results were analyzed by the ΔCt method and the 18S gene was used to normalize the corresponding Ct values [24]. Primers used in this study are listed in supplementary data (Table S33). Twenty-five days post CLCuD inoculation, the leaf discs from the top three leaves of each whitefly exposed and grafted cotton plants were excised and pooled for CTAB-mediated DNA extraction. The purified DNA was nanodrop and diluted to a final concentration of 10 ng/µl. For qPCR analysis and standard preparation previously optimized protocol by Shafiq et al. [49] was followed. The dilutions of plasmids were made in the range from 2 ng, 0.2 ng, 0.02 ng to 0.002 ng for standard curve preparations.

## RESULTS

### Genome-wide identification

The whole-genome screening analysis identified a total of 12,822 NBS encoding genes across 34 species. The species-based wide-genome identification demonstrated that the number of genes present in each genome was independent of its genome size. As the genome size to no. of NBS gene regression (R^2^) value was observed as belove the cut of value (R^2^ = 0.015). Interestingly, we observed a correction of genome-size to no. of NBS genes in *Gosypium sp* (R^2^=0.669). For instance *Gossypium hirsutum* with genome size 2.4Gb has highest no. of NBS genes (708 NBS genes), followed by *Gossypium barbadense* (622 NBS genes) with genome size 2.3Gb, and *Gossypium arboreum*, a diploid genome with 1.7Gb genome, has 365 NBS genes. Similarly, the *Gossypium raimondii* with 0.7Gb genome has 323 NBS genes. The detail plant species and number of identified genes were given in the supplementary files. (Figure 1, Table S1, Table S2).

**Figure 1.**
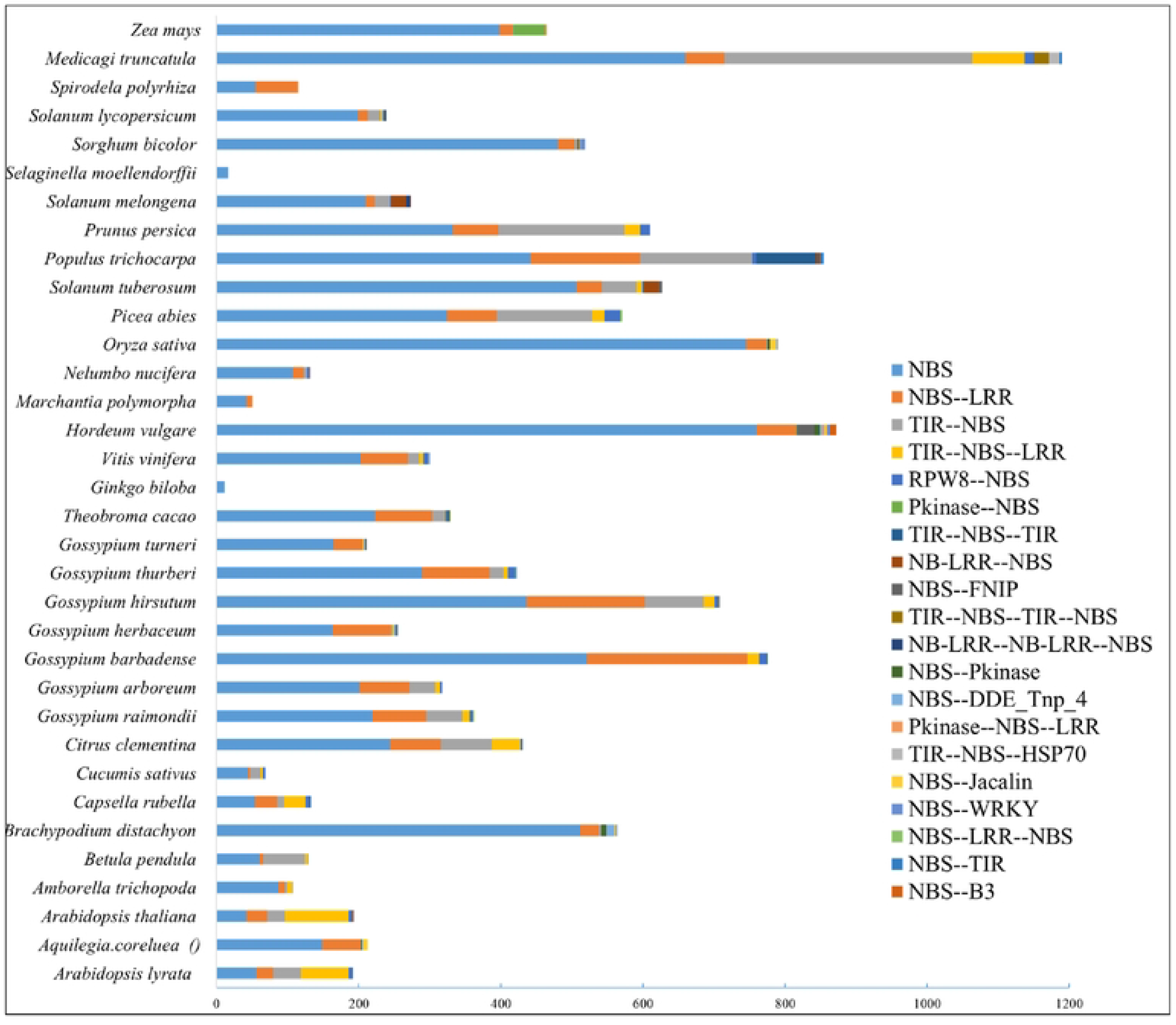
Genome-wide identification and classification of NBS genes in Land plants. The length of the bar represents total number of genes and different colors demonstrate top 20 subclasses, based on domain architecture.

### Classification of NBS genes in land plants

The *Pfam* domain analysis identified several decoy domains in addition to the NBS domain. The deep analysis of domain architecture reported several functional protein domains associated with NBS domains (Table S2). So, based on the conserved domains, motifs, and their sites in the primary sequences of NBS proteins, all 12822 genes were divided into 168 classes with respect to the presence /absence and copy number of associated domains (Table S3). Of these classes, the 33^th^ class (only NBS domain) has 8,591 NBS genes, showing largest class of NBS superfamily and most of the classes were highly specific with few number of NBS genes. So, based on the observation, it can be estimated that the main superfamily of NBS proteins contain only NBS domain. The species-wise distribution and diversity of identified NBS classes were also interesting. As the lower plants possessed only either one or two specific classes e.g., *Selaginella moellendorffii* belong to a lycophyte, and possessed only 33^th^ class (only NBS single domain) with 16 NBS genes. Similarly, *Spirodela polyrhiza* is a species of duckweed known by the common name common duckmeat, has only two classes 33^rd^ and 60^th^ (NBS and NBS-LRR). In contrast, the evolutionary development and adaptation process causes drastic changes in the genome of higher land plants. As moved from simple to complex plants, the NBS classes became more complex. For instance, *Hordeum vulgare*, barley, a member of the grass family, is a major cereal grain grown in temperate climates globally and has the highest number of NBS classes several unique NBS domain architectures. Similarly, *Vitis vinifera*, the common grape vine, a species of flowering plant, native to the Mediterranean region, also has twenty NBS classes. Among all land plants, the most common domain architectures were detected as NBS, NBS-LRR, TIR-NBS, TIR-NBS— LRR, and RPW8—NBS domain architectures. However, we also observed highly species-specific classes like Methyltransf_11—NBS, NBS-Glyco_transf_8, TIR-NBS-Lectin_legB-Pkinase, TIR-NBS-RHD3 and NB-LRR-NB-LRR-NBS-Retrotran_gag_2. Overall, out of 168 classes, 94 classes were specific-specific, and some classes were family specific and were genus specific like the Gossypium species showed a close relationship regarding the classes. The tetraploid species (*G. barbadence* and *G. hirsutum*) of Gossypium have more diversity rather than other diploid species (Figure 1, Tables S3, Table S4).

### Orthologs analysis: orthogrouping, duplication and overlapping

The orthologs groping analysis of 12,822 NBS in 33 land plants demonstrated that 91.8% of total NBS proteins were assigned orthogroups. However, only 8.2% of NBS proteins were highly unique and could not fit into any orthogroups. A total of 664 orthogroups were predicted and out of these, 347 orthogroups (holding 1135 NBS proteins) were species-specific orthogroups and the mean and median orthogroup size was 18 and 3 genes, respectively (Table S5). The species-based comparative genomics overexposed the genetic makeup of different land plants regarding NBS genes. We identified many common/core genes as well as species-specific unique genes based on sequence divergence. For instance, in the case of *Arabidopsis layrata*, a total of 209 NBS genes were predicted and categorized into 37 orthogroups (with 2 species specific orthogroups) covering 206 NBS genes. The remaining three genes did not fit any orthogroups due to their clear sequence divergence as compared to other NBS genes. Similar results were also observed for other land plants. For instance, in the *Gossypium sp*, species i.e., *Gar, Gba, Gbi, Ghe, Got, Gra* did not show any species-specific unique orthogroup or ortholog. But observed some unique NBS genes which were categorized as “unassigned orthogroup genes” (Table S6). The number of genes in orthogroups of each species was also assessed and identified that the “OG_000_” orthogroup possessed the highest number with 2535 NBS proteins and shared all species except some lower plants including *Gbi, Mpo*, and Smo. In addition, some orthogroups were highly specific to species like OG_400_ was only present in *Al* with *ALNBS134 and ALNBS135* genes. Similarly, the OG_024_ was specific to Medicago with more than 15 NBS genes and the OG_454_ (*GhirNBS444* and *GhirNBS445*), OG_455_ (*GhirNBS626* and *GhirNBS627*) was specific to *G. hirsutum.* Similar unique genes were also identified in other land plants (Table S7, Table S8, Table S9, Table S9, Table S10). The orthogroups duplication event analysis conformed several duplication events during the evolutionary process e.g. the OG_000_ duplicated 1932 times with 100% confidence and 1600 times with a 50% confidence level. Similarly, other orthogroups were also passed by some duplication events (Table S11, Table S12).

In order to find the relationships among different species the orthogroups and orthologs overlapping were also assessed and compiled into clusters using tendogram. The result demonstrated that *OS, Hvu, Zm, Bra,* and *Sobic* formed one clade, and *Sopyc, pgsc, sme* form another clade. Similarly, the Gossypium sp. Including *Got, Gtu, Gra, Gba, Ghi, Gar*, and Ghe formed one clade. In addition, the Gar (A-genome) and Ghe (A2-genome) shared sister branched and the Ghir (AD-genome), Gbar (AD2-genome) formed another sister branches showing the highest genome similarity index (Figure S1, Figure S2, Table S13).

### Phylogenetic Tree Analysis: species-base, gene-based and terminal duplication events

The species-based phylogenetic tree and duplication events at internal and terminal nodes indicated the evolutionary relationship of NBS genes in land plants. A close relationship was observed among different species of belong to the same genus e.g. *Solanum* genus forms a separate clade covering *S. tuberosum, S. lycopersicum* and *S. melongena* and all *Gossypium* species shared a single clade with the most conserved genome of *G. raimondii* and a least common ancestor with *T. cacao*. Similarly, the *N. nucifera* act as the least common ancestor of monocot flowering plants including *Z. mays, O. sativa, S. bicolor and H. vulgare*. The duplication events at terminal nodes demonstrated that *G. biloba* was highly conserved and did not demonstrate any duplication at the terminals as compared to other land plants. Entering into the *Gossypium* genus, the N_24_ node further duplicated more than 210 times and separated other *Gossypium* species from *G. raimondii* (Table S14). In conclusion, the species-based phylogenetic tree and gene duplication events at internal and terminal nodes demonstrated the expansion of NBS gene family from lower plants to higher plants. The duplication events also cause the gain and loss of NBS-associated terminal domains, as we observed in the domain architecture analysis. This diversity in NBS proteins might be an adaptation of land plants (Figure 2)

**Figure 2.**
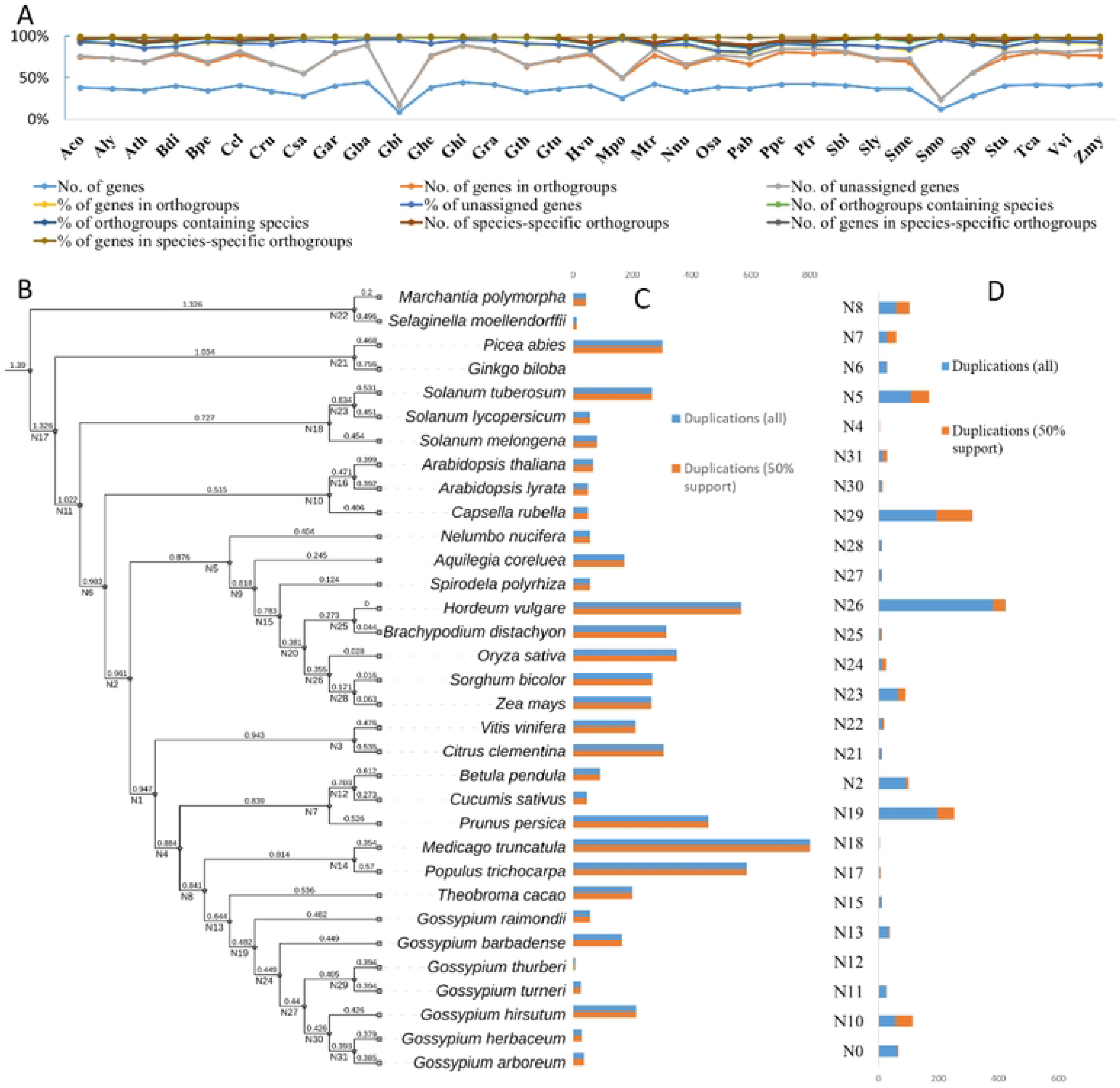
Evolutionary study of NBS genes in land plants. A) Basic statistics of orthogroups with abbreviated species na1ne. B) Species-based phylogenetic tree, N represents internal nodes in phylogenetic tree. C) Gene duplication events at tenninal nodes of the phylogenetic tree, and D) gene duplication events at internal nodes of a phylogenetic tree.

The gene-based phylogenetic tree was divided into 282 sub-clusters and each cluster was correspondence to the orthogroups. The highest number of genes was observed in cluster OG_000,_ which gradually decreased to and decreased to OG_282._ In addition, some clades were highly species-specific and some were genus while very few were family specific. Due to the large number of genes in top orthogroups only OG2 was present in the phylogenetic tree (Figure S3)

### RNA-bases expression profiling

For a deep understanding of NBS genes in different tissues under different biotic and abiotic stresses, we have selected three plant species including *A. thaliana, Z. mays, G. arboreum* and *G. hirsutum*. Based on the RNA-seq expression data most important putative genes were further filtered for detail study. The tissues specific expression profiling of NBS genes in *A. thaliana* generally demonstrated that most of the genes are differentially expressed in leaf, shoot, seedling, flower, and silique. However, very little or negligible expression observe in pollen, endosperm, embryo, and seed. At orthogroups (OGs) level, some OGs were highly specific to some tissues like OG_6_ (*At4G33300_OG6_* and *At5G04720* _OG6_), OG_15_ (*AT5G45490* _OG15_) had the highest expression in root tissue (Table S15). Similarly, the OG_2_ (*AT1G61300*, *At5G63020*, and *AT1G61190*) showed similar patterns in leaf, shoot, and seedling. The cladogram among tissues and genes also demonstrated a co-expression network of genes. In the case of Zmy tissues, three major co-expression pattern network was observed among NBS genes.(Figure 3, Table S16, Table S19, Table S22, Table S25).

**Figure 3.**
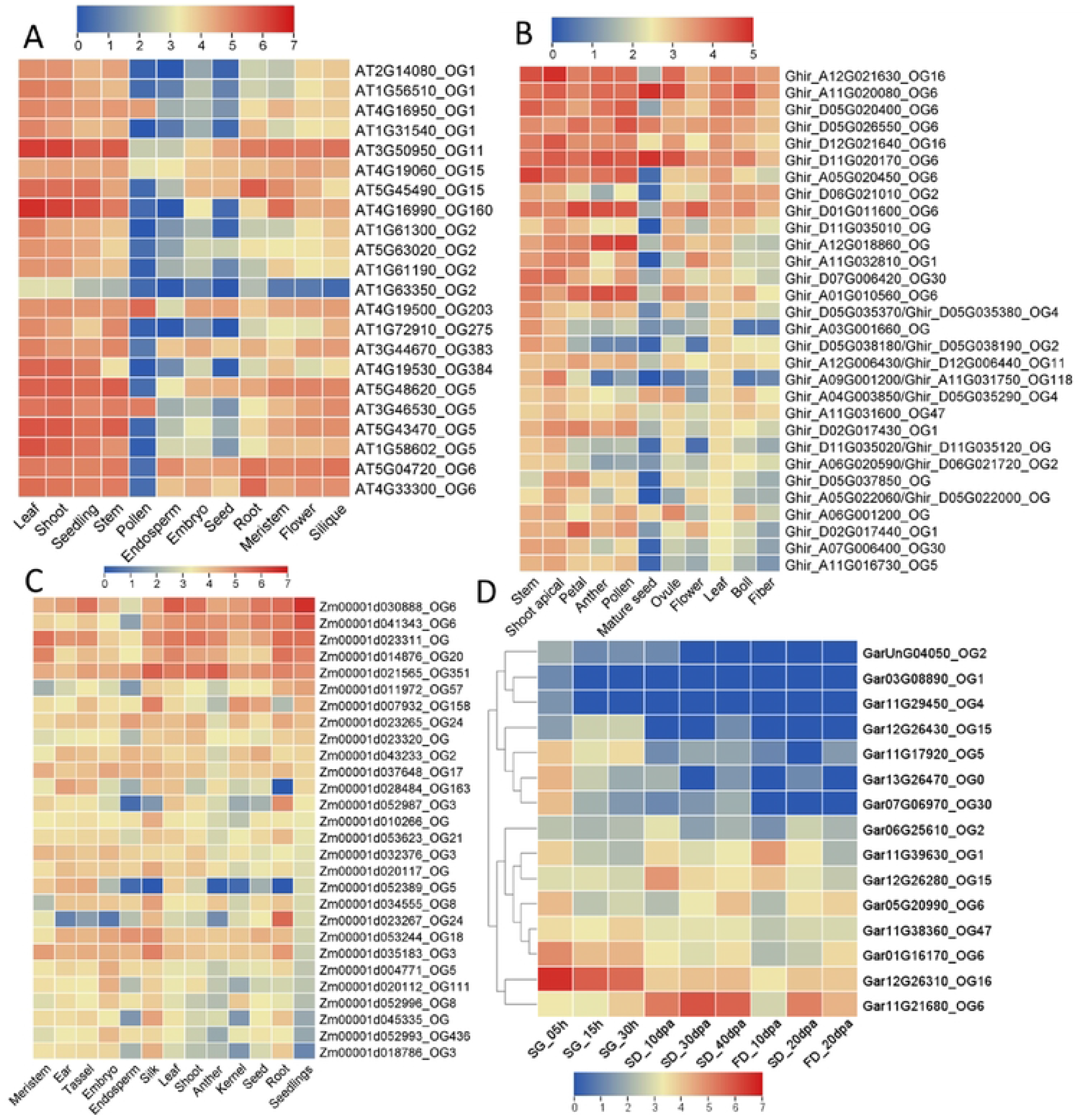
RNA-seq based expression profiling of NBS gene in different tissues. The OGs associated with gene IDs represent the Orthogroups. A) *A. thaliana,* B) *hirsutum* and C) *Z. mays* and D) *G. arboreum*

**Figure 4.**
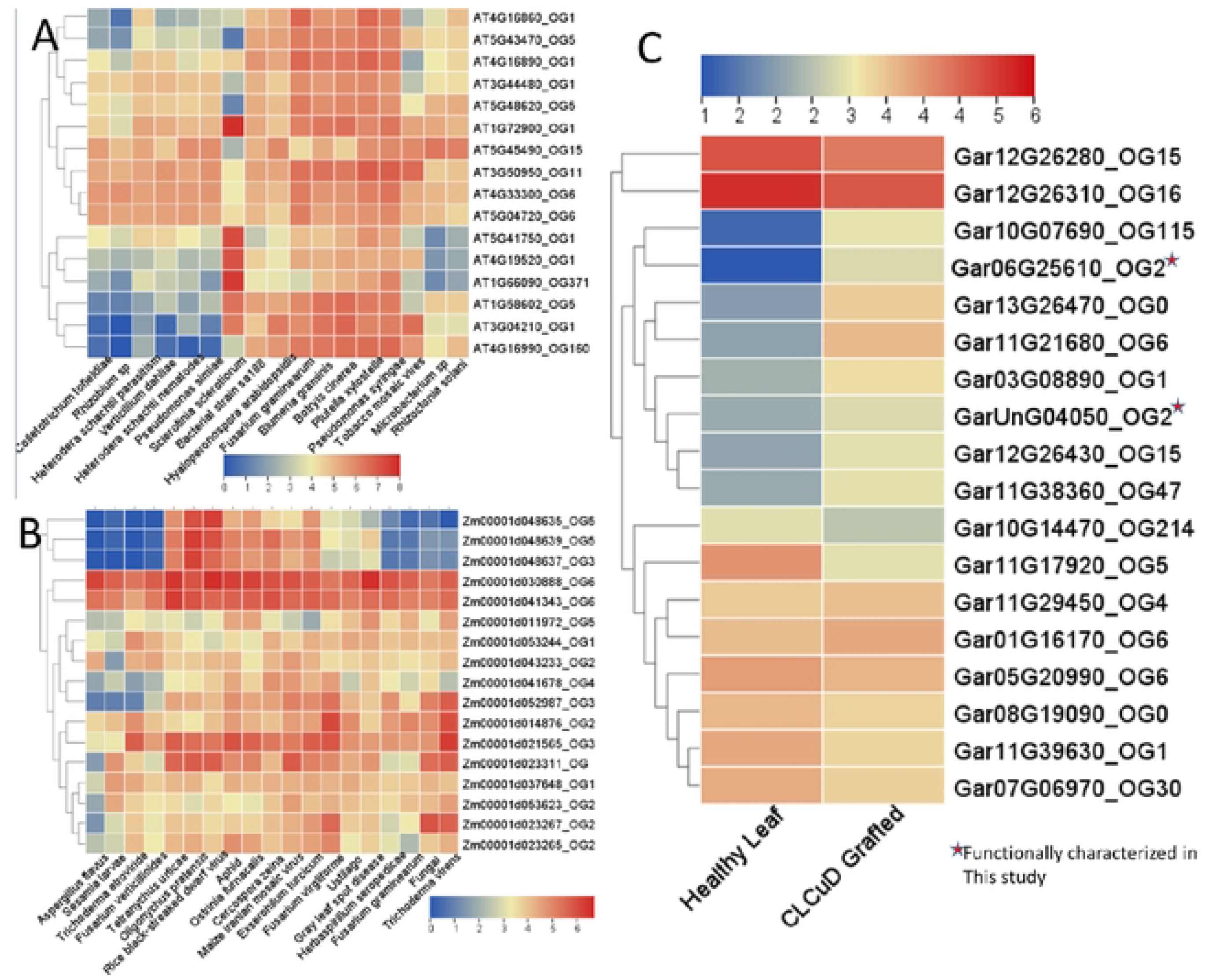
RNA-seq based expression profiling of NBS gene under different biotic stresses. The OGs associated with gene IDs represent the Orthogroups. A) *A. thaliana,* B) *Z. mays* and C) *G. arboreum*

The RNA-seq data of abiotic stresses included Oxidative stress, UV, Nutrient deficiency, dark, salt, cold, drought, heat, ozone, and wounding stresses. Generally, it was observed that the OGs members mostly co-expressed under various abiotic stresses. In *A. thaliana*, the NBS genes are most commonly expressed in all stresses except oxidative and UV stresses. Among OGs, the OG_6_, OG_11,_ and OG_1_ showed similar expression patterns with the highest expression under most of the stresses. Similarly, the OG_6_, OG_0,_ and OG_351_ formed one co-expression clade in *Z. mays* plant under various abiotic stresses (Figure S4, Table S17, Table S20, Table S23).

The biotic stress expression profiling included different pathogens covering from aphids to viruses (aphids, nematodes, fungi, bacteria, and viruses). The analyses identified several NBS gene responses under different pathogenic stresses in *Arabidopsis, Z. mays, G. arboreum* and *G. hirsutum*. For instance, the OG_1_ (*AT1G66090_OG371_, AT4G19520, AT5G41750, and AT1G72900*) significantly upregulated under *Slerotinia sclerotiorum*. Similarly, under viral disease the OG1, OG5, and OG6 showed putative responses during viral infection in Arabidopsis and *Z. mays*. The *G. arboreum* is naturally resistant to several viral diseases, so we have taken the CLCuD grafted RNA-seq for the identification of NBS genes role in the presence of viruses. We identified several differentially expressed NBS genes in *G. arboreum* under grafted-CLCuD. The OG2 (Gar06G25610, GarUnG04050), OG6 (Gar11G21680), OG4 (Gar11G229450) and OG115

(Gar10G07690) showed significantly upregulation under CLCuD. The Gar06G25610_OG2_ was further validated through gene silencing approaches in this study (will discuss later) (Figure S5).. As we are so interested in the identification of most putative NBS genes and their role in *G. hirsutum* under cotton leaf curl disease, which is Sone of a major challenges in Pakistan, we presented a two-contrast agronomic trait associated accession (Mac7 (a highly CLCuD tolerant *G. hirsutum* accession developed by USDA, but low production) and Coker312 (highly susceptible, most regenerative accession, commonly used for tissues culturing, also posseted important agronomic traits) RNA-seq data. The results were surprising, we found differential regulation of NBS genes in tolerant accession (Figure 5, Table S18, Table S21, Table S24, Table S26). In summary, we found that the OG_0_, OG_2_, OG_5_, OG_6,_ and OG_15_ have a highly responsive role in different tissues and under various biotic and abiotic stresses in *Arabidopsis, Z. mays*, *G. arboreum* and *G. hirsutum*.

**Figure 5.**
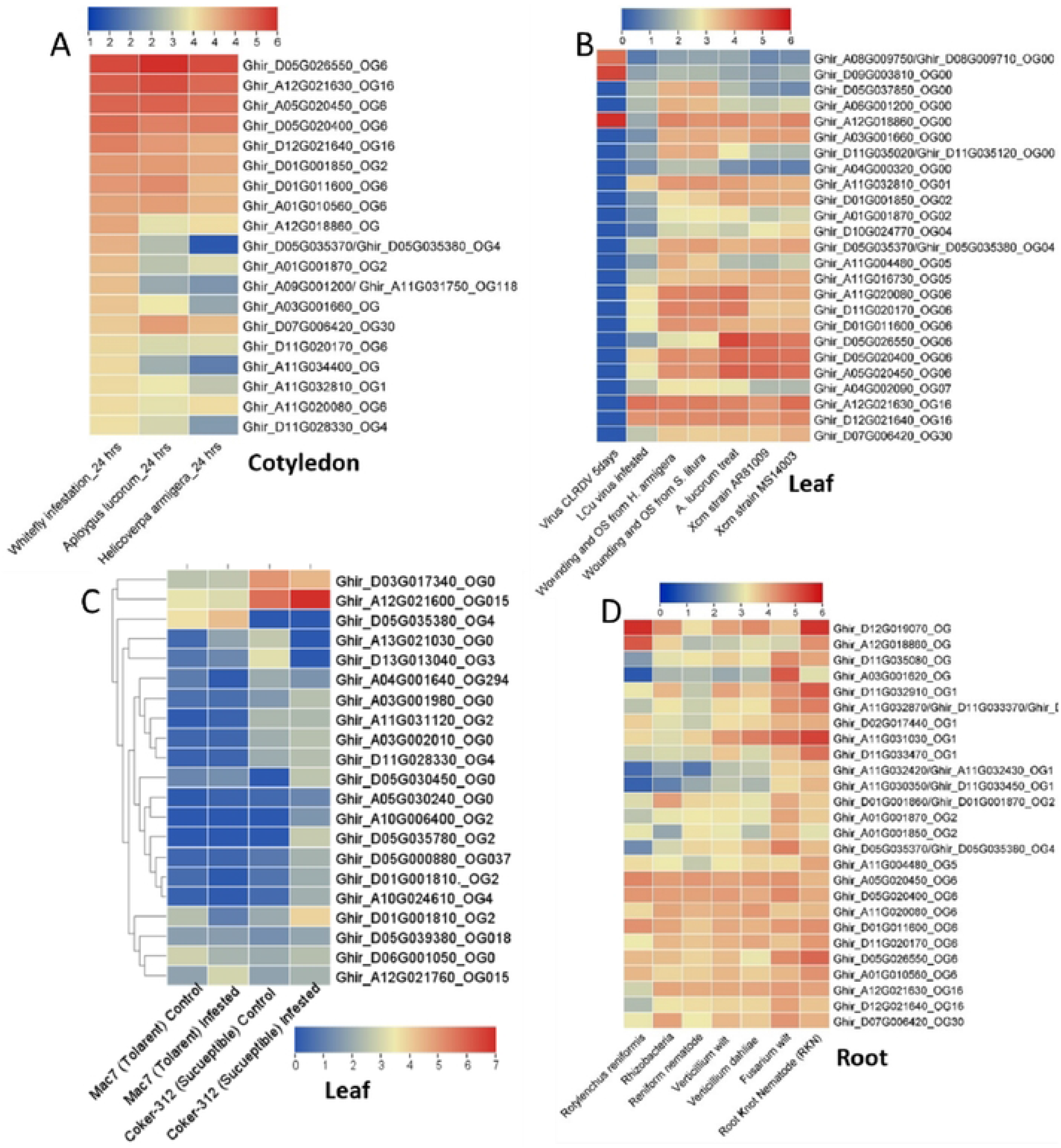
Expression profiling of G. hirsutu1n NBS genes in different tissues under various biotic stresses. A) Reference genome RNA seq data in cotyledon tissue, C) expression of NBS gene in two contrasting accessions, I; Mac7 (tolerant to CLCuD), 2; Coker3 I2 (highly susceptible to CLCuD) under CLCuD infection in leaf. D) under various root-associated pathogens in root tissue.

### Variants detection in tolerant and susceptible accessions

The genome-wide genetic diversity of Mac7 (*G. hirsutum* tolerant) and Coker312 (*G. hirsutum susceptible*) with reference to TM-1 *G. hirsutum* reference genome, identified several SNPs and InDels in NBS genomics regions of Mac7 (InDels: 2989, SNPs: 3594) and Coker 312 (InDels: 2646, SNPs: 2527). The identified variants were characterized into four impact levels *i.e.* impact as high (affecting splice-sites, stop and start codons), moderate (non-synonymous), low (synonymous coding/start/stop, start gained), and modifier (upstream, downstream, intergenic, UTR). A comparative study of high-impact SNPs and InDels associated genes between Mac7 and Coker identified several unique and common variants e.g., Coker 312 (7 SNPs, 10 InDels) and Mac 7 (22 SNPs, 26 InDels) unique variants. While 4 genes were common between the two accessions. Similarly, the modifier and moderate and low-impact variants also observed in the two accessions. See the supplementary file for a detail comparison and associated genes (Figure 6, Table S27, Table S28, Table S29).

**Figure 6.**
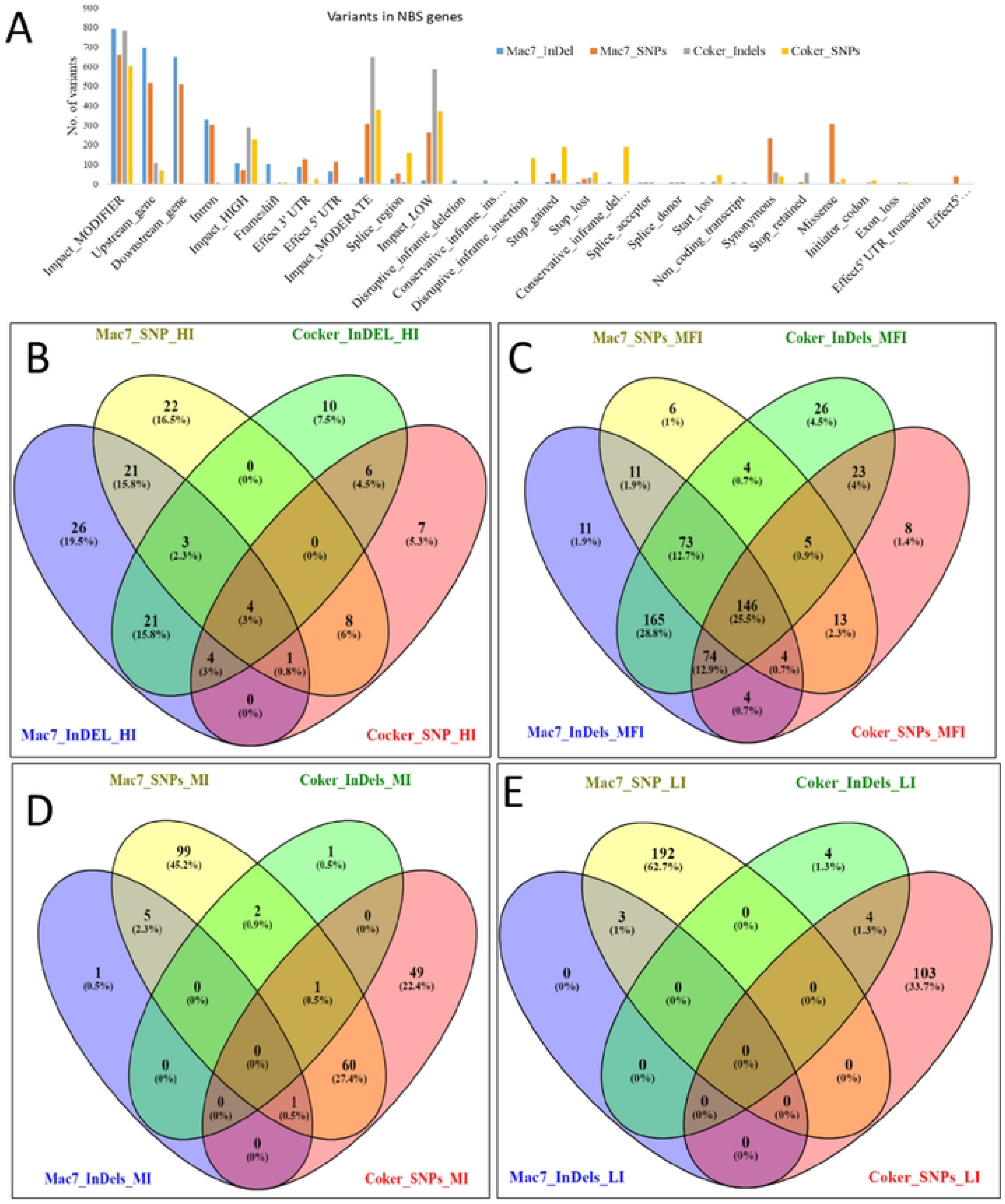
Genetic variations of NBS genes between Mac7 and Coker 312.A) variants distribution region Based on i1npact level of variants on gene functions. A) Hl; High Impact, B) MF!; Modifier Impact, C) MI; Moderate Impact, D) LI; Low Impact, comparison at SNPs(single nucleotide polymorphisms) and InDels (Insertion and Deletion)

### Physiochemistry, gene ontology and metabolic pathways

The amino acid (AA) sequence analysis of *Gossypium hirsutum* showed a length ranging from 500-1500 amino acid (AA), molecular weight (MW) ranged from 50-150 kDa, isoelectric point (pI) value ranged from 2 to 6, charge ranged from +55.5 to −44.5 and the Grand Average of Hydropathy ranged from 0 to −0.2. The gene ontology comprises three components; biological process, molecular function, and cellular components. In the case of molecular function, the results demonstrated that the NBS is mainly involved in the Adenosine diphosphate (ADP) and Adenosine triphosphate (ATP) binding activity with signal transduction. The KEGG pathway analysis clearly indicated the role of the NBS gene in plant-pathogen interaction (PPI) and signaling pathways (SP). The promoter analysis identified several stress-responsive elements in the promoter region of NSB genes. A total of 72 cis-regulatory elements were identified some of the responsive elements were present in all NSB genes like TATA-box, CAAT-box, Box, ARE, G-box, GT1-motif, ABRE, CGTCA-motif, TGACG-motif, TCT-motif, TCA-element, MBS, GATA-motif, CAT-box, and O2-site. Most of the cis-regulatory elements belonged to stress responses (Figure S5, Table S30).

### Molecular docking, and proteins-protein interaction

For the proteins modeling, we selected genes based on upregulated in *G. arboreum* (Naturally resistant to CLCuD) under grafted-CLCuD and whitefly induced in Mac7 (*G. hirsutum*, tolerant to CLCuD) covering OG_0_, OG_2_, OG_15_, and OG_43_.The selected genes’ translated proteins were used for 3D structure prediction using I-TASSER server. The PDB database proteins 6J5T-C,6S2P-N, 7CRB-A,4TZH-A, and 4U09-A were used as template sequence. The 3D molecule of ATP and ADP were downloaded from cheminformatics database. The docking results demonstrated a stable interaction of NBS genes with ATP and ADP with a range of −7Kcal/mol to 8.2kcal/mol, except *Gar09G25760_OG43* and *Ghir_A12G021600_OG15*, which showed below −6.8 Kcal/mol and −6.6Kcal/mol, respectively. The binding affinity of *Gar12G23120_OG0* showed that it has a more stable interaction with ATP as compared to the ADP molecule. Similarly, the OG2 genes (*Gar01G01860, Gar06G24920*) showed more affinity to ATP molecule as compared to ADP molecule. In contrast to this, the OG0 (*Gar11G29700, Ghir_D13G021900*) showed more stable interaction with the ADP molecule. The interacting residues of NBS proteins varied from ATP to ADP e.g. the ATP binds with Lys179, His240, Trp139, Val261, Gln 264, Glu142, and Val 284, whereas the ADP binds with Asp143, Glu142, Asn289 and Lys286 in Gar12G23120_OG0 protein. Similarly, other binding results also demonstrated similar patterns (Figure S6, Table S31). For further detailed molecular interaction of OG2 genes (*Gar06G24920_OG2*) with CLCuD viral proteins (AC1, AC2, AC3, AC4, AV1, and AV2), we assessed the interaction level by Gibs free energy. The *Gar06G24920_OG2* protein demonstrated the highest stability value (−1352Kcal/mol) with AC1 viral proteins and followed by AV2 (−1103.8), AV1 (−1026.4) and so on. Similarly, another member of OG2 group protein (*Gar06G25610/Gohir.A06G19220/ Ghir_A06G020580*), also demonstrated high interaction value with AC1 (−1506.6 Kcal/mol) and AV1(−1255.8 Kcal/mol), AC3 (−1230.8Kcal/mol). The active residues were also changed with the viral protein complexes. (Figure S7, Table S32)

### Silencing of OG2 (G2) genes enhanced virus titer in *G. arboreum*

TRV-based VIGS assay in cotton is a well-established approach to conducting functional studies of OG2 (*Gar06G24920_OG2) genes*. FDH-228 plants were inoculated with VIGS constructs to silence the subjected genes, 15 days post infiltration, a completely bleached phenotype was visible on TRV-GrCLA1 inoculated cotton plants, at this time, RT-qPCR based gene silencing was performed (Figure S8-S9). The results demonstrated a considerable reduction in the expression level of the G2 gene in FDH-228 silenced plants, compared to the TRV:00 inoculated control plants (Figure 7A). The graft-mediated CLCuD inoculation has been well demonstrated by Ullah et al. [60] in *G. arboreum* plants. Since this breakthrough, it was possible to identify the resistance imparting genes of *G. arboreum* against CLCuD. After assessing the gene silencing, an equal number of silenced and control plants were inoculated with CLCuD by grafting CLCuD harboring scions and exposing them to viruliferous whitefly. 25 days post-CLCuD exposure, enhanced virus concentration was witnessed in G2 (*Gar06G24920_OG2*) silenced plants in comparison to TRV:00 inoculated controls both under graft and viruliferous whitefly exposure (Figure 7B). Minor disease symptoms were evident only on graft-inoculated G2 (*Gar06G24920_OG2*) silenced plants (Figure 7C-G). These results suggested the likely involvement of the G2 (*Gar06G24920_OG2*) gene in CLCuD resistance in FDH-228 plants.

**Figure 7.**
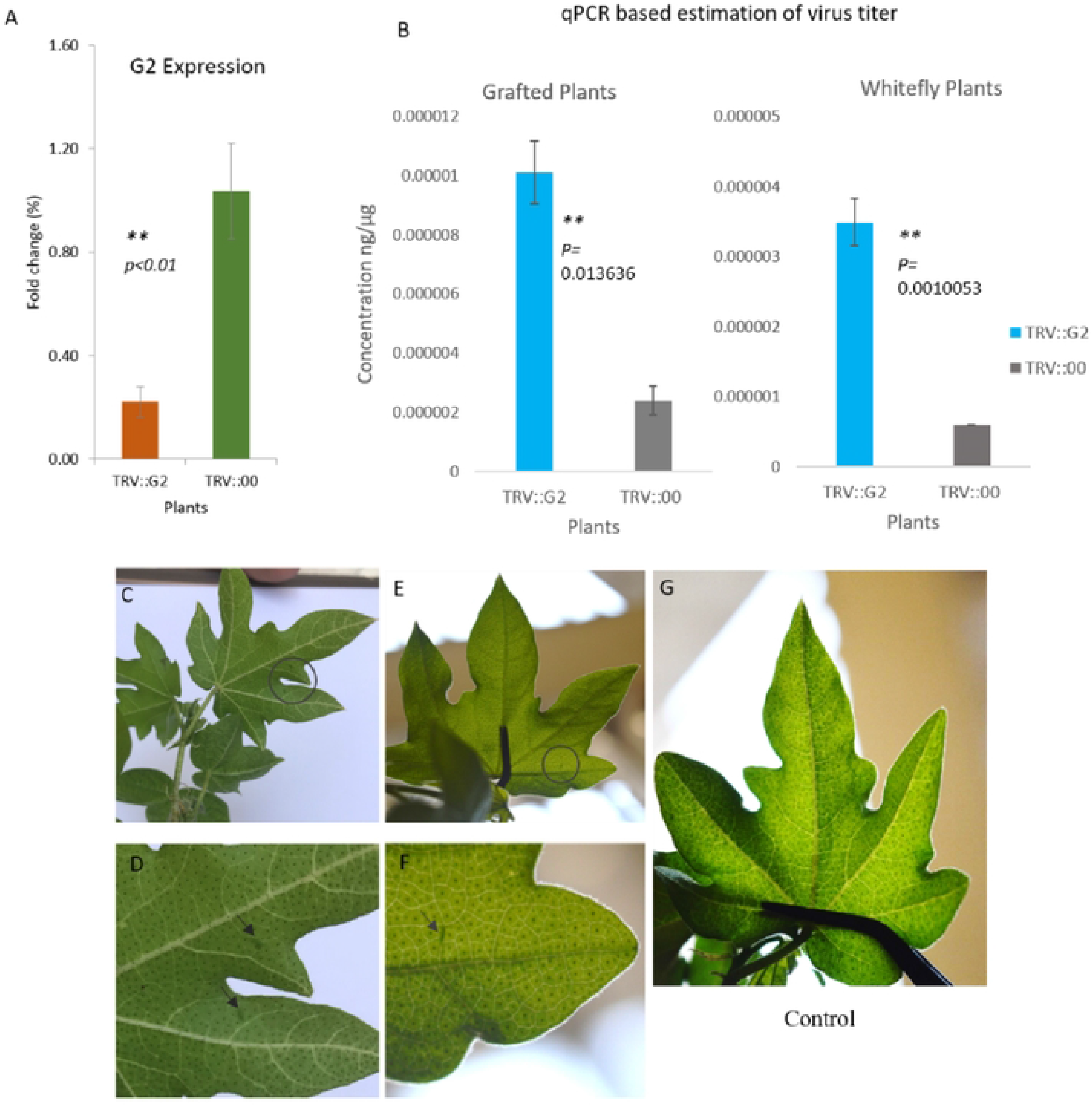
qPCR-based estimation of gene expression and CLCuD Symptoms development on VIGS plants. Panel (A), Orange is representing decreased expression of G2 gene in *G. arboreum* plants. Green is showing expression of G2 in empty vector inoculated control plants. Typical CLCuD symptoms of lower grade appeared on G2 VIG$ plants. Panels (C, D) and (E, F) are showing minor vein thickening on G2 VIGS plants G is presenting TRV:00 inoculated FDH-228 leaf with no symptoms. Bis showing qPCR-based estimation of virus titter in both graft and whitefly exposed VIGS plants of *G. arboreum.* ** is for a significant difference.

## Discussion

In the last two decades, several types of resistance (R) genes have been discovered. The majority of R genes found so far are members of the nucleotide-binding site (NBS)-leucine-rich repeat (LRR) receptor (NBS-LRR, also known as NLR) gene family [50]. NLR genes have been found in plants and their origin may be traced back to the common ancestor of all green plants. Prior to the split of green plants, phylogenetic studies revealed that NLR genes had diverged into separate subclasses [51, 52]. According to genome-wide studies, different species of plants possess a varying number of NLR genes. For example, in the Poaceae family, the number of genes ranges from 145 (maize; a diploid plant) to 2298 (T*riticum aestivum*; a hexaploid) [53]. Our genome-wide identification result also identified a various number of NBS genes in a land plant covering mosses to angiosperm [54]. Furthermore, we have also observed that some of the green plant ancestors like *Coccomyxa subellipsodiea, Klebsormidium flaccidium, Micromonas pusilla, Physcomitrella Patens,* and *Volvox carteri* do not possess NBS genes, indicating the lack of such resistance gene in ancestor plants. NLR gene expansion and contraction can support the “birth-and-death hypothesis” that predicts fast NLR evolution in land plants, might be due to duplication events at the genome, and segmental levels [54, 55].

NLR genes are frequently clustered in complex clusters, a structure that may favor NLR dynamic development and diversification to cope with rapidly evolving pathogens [56]. The detection of certain pathogen effectors activates NLRs, resulting in a robust immune response that is frequently linked with localized programmed cell death, known as the hypersensitive response (HR) [57]. Most NLRs feature an N-terminal extension consisting of a Toll/interleukin-1 receptor (TIR) domain, a coiled-coil domain (CC), or a divergent coiled-coil domain (CCR) identical to the Resistance to powdery mildew 8 (RPW8) domain [58]. Generally, there are three kinds of NLRs based on the N-terminal domain and evolutionary history of the NB-ARC which are 1) TIR-NLRs (TNLs), 2) CC-NLRs (CNLs), and 3) RPW8-NLRs (RNLs) [12, 59]. Our classification system was unique and identified several N-terminal and C-terminal helper domains, in addition to NBS domains, e.g. Zf-BED, WRKY, BRX, LIM, DA1-like, Plant_tran, zf-RVT, Myb_DNA-binding, AP2, DDE_Tnp_4, Thioredoxin, Jacalin, zf-BED, etc. These additional domains are known as immunological receptors with integrated domains (IDs) that resemble pathogen targets and are activated in response to effector change [60]. Previous literature data divided NLRs into two functional groups: 1) direct/indirect sensor NLRs that detect invasion and 2) helper NLRs that are genetically necessary for immunological activation by other NLRs [61, 62]. Based on the presence and absence of the 2^nd^ group, we have classified all 12,822 NBS genes into 168 classes. The number of different classes in different plant species also gives genetic features of land plants and their evolutions. Some of these identified classes are reported in the literature and well-characterized. As the RNL clade is usually characterized by a low copy number [63, 64], with the exception of Gymnosperms (there are 31 ADR homologs in spruce) [65]. RNLs exhibit extraordinary intron conservation in *Amborella* and dicots, which share four introns, but monocots have three introns (the second is missing) [66, 67]. NRG and ADR were separated before angiosperms diverged and they are still retained in syntenic blocks across flowering plants and several lineages have lost NRG genes but not ADR genes [68]. The TNLs are divided into two subfamilies: 1) TIR1 and 2) TIR2 [69], while in some plants such as monocots, only TIR2 NLRs are maintained [69, 70]. Many dicot species have TIR1 NLRs, however, other dicot lineages are devoid of them [71]. So, the presence and absence of these newly identified classes in our study further required functional characterization.

The evolutionary study of NBS genes in 33 land plants provided several common patterns of NLR evolution that were not visible in studies of a single species or plant family. A total of 664 orthogroups (OGs) were predicted among the land plants with only 8.2% of the total genes with unique sequences. Of these, we found many core and conserved OGs in plant species-specific lineage. The OG00, was shared with all plants except some lower plants, showing orthologs reduction due to their small and simple genome. The OG2 was another orthogroups that was found as the core orthogroups among land plants. The genome and tandem duplication events also cause the extension of OGs in the genome and cause genomic diversity [72, 73]. The diversity analysis of NBS at the genic, genomics, and species level also demonstrated duplication events at internal and terminal nodes of the species-based phylogenetic tree during the evolutionary process.

The RNA-seq-based expression profiling of the NBS gene in *A. thaliana*, *Z. mays*, *G. hirsutum* and *G. arboreum* demonstrate the significant role of the NBS gene in plant growth and development [74, 75]. As in the control of NLR activity, gene transcription is an early regulatory step. To acquire resistance, proper NLR gene transcription is necessary as excessive transcription can cause programmed cell death, which is detrimental to plant growth and development. Overall expression profiling suggested that the OG2, OG6, and OG15 members showed the highest putative role in different tissues, under various biotic and abiotic stresses. The highest expression of these genes in the given plant species under biotic and abiotic stresses might be due to the presence of stress-responsive cis-regulatory elements in the promoter region, which was clearly observed in the promoter sequences of *G. hirsutum* e.g. Box, ARE, G-box, GT1-motif, ABRE, CGTCA-motif, TGACG-motif, TCT-motif, TCA-element, MBS, GATA-motif, CAT-box, O2-site. In addition to the presence/absence of stress-responsive elements, epigenetic markers such as histone post-translational modifications and DNA methylation can also influence NLR gene expression by modulating their chromatin structure. NLR gene expression may be influenced by DNA methylation at the promoter, as the methylation in the promoter of the *Arabidopsis* TNL gene (MG1), upregulated flg22 treatment, implying dynamic DNA methylation in NLR promoters during biotic stress. Additionally, the genetic variation in the UTRs, splicing sites, and exonic regions of NBS genes in the susceptible and tolerant *G. hirsutum* also depicted their role in the two lines under cotton leaf curl disease.

For long term strategies to control disease in cotton and other important crops, the deployment of genomics tools including, NGS sequencing, exome sequencing, and genotype by sequencing, are helping to identify genetic variations (InDele, SNPs, SSR) associated with the functional diversity, which is further in marker-assisted selection (MAS) approach during breading of wild (resistant) and cultivars (susceptible) [37, 55]. In the current study, we have made a comparison of genetic variation between a highly virus-resistant *G. hirsutum* accession and a highly susceptible cultivar and found several SNPs and InDels in tolerant accession. As breeding of virus-resistant cotton varieties with sufficient genetic diversity has been suggested as a durable strategy for controlling the disease effectively [76, 77] and the genetic basis of resistance and its inheritance are the key components for designing breeding strategies. In cotton plants, very little is known about the genetic variants associated with cotton leaf curl disease resistance. Therefore, our identified genetic variants might help develop CLCuD-resistant varieties, which is our next project.

Plant NLR proteins have been discovered to have a central NBS domain that resembles the AAA-ATPase family [78, 79]. The ADP and ATP are the two molecules that have a putative role in the activation and deactivation of NBS domain [80]. The binding of ADP with NBS domain, cause deactivation, while its exchange with ATP opens the active site of NBS domain and activate the molecular mechanism [81]. After activation, two intracellular proteins CC-NBS-LRR and TIR-NBS-LRR recognize viral effector molecules to activate the defense responses [82]. So, in order to undertrained the deep mechanism of NBS protein activation in cotton plants, we have presented a comparative molecular interaction of ADP and ATP with NBS proteins and found that the OG2 genes (*Gar01G01860, Gar06G24920*) showed more affinity to ATP molecule as compared to ADP molecule, suggesting their active role in defense response. Furthermore, the functional validation of OG2 genes (*Gar01G01860, Gar06G24920*), in *G. arboreum* (a naturally immune land race) through virus-inducing gene silencing also increased the viral disease system in silenced plants as compared to control plants.

## Conclusion

The identification and analysis of resistance genes in plants have revealed interesting patterns of their evolution and diversification. The majority of these genes belong to the NBS-LRR receptor gene family, which can be traced back to the common ancestor of all green plants. The number and type of NLR genes vary greatly among different species, indicating their adaptation to cope with rapidly evolving pathogens. NLRs are frequently clustered in complex clusters, which may facilitate their diversification and development. Different classes of NLRs have been identified based on their N-terminal and C-terminal helper domains, some of which resemble pathogen targets and are activated in response to effector change. While the study of NBS genes in different plant species has revealed common patterns of evolution, the presence and absence of newly identified classes require further functional characterization. Furthermore, the NBS gene plays a significant role in plant growth and development, as demonstrated by RNA-seq expression profiling in various plant species. Proper transcription of the NLR gene is necessary for acquiring resistance, as excessive transcription can cause programmed cell death that is harmful to plant growth. The OG2, OG6, and OG15 members showed the highest putative role in different tissues, under various biotic and abiotic stresses, possibly due to stress-responsive cis-regulatory elements in the promoter region. Finally, the functional validation of OG_2_ in resistant cotton plant demonstrated its putative role in plant resistance during cotton leaf curl disease infection. Overall, these findings provide insights into the genetic features and evolution of land plants.

## Acknowledgments

We are thankful to Bioinformatics Group, Molecular Virology and Gene Silencing Lab, National Institute for Biotechnology and Genetic Engineering (NIBGE), and the University of Management and Technology (UMT), for providing suitable research facilities and a conductive environment for this work.

## Author contributions

SM and IM supervised this work. AH and MQA designed this work and wrote the first draft. AH, AAK, NZ, MF, MAM, AA, and AN performed the How Bioinformatics. MQA performed the gene silencing experiment. MS, SM, HUR and IM funded this work. MSM, HUR, MF, HUR, SM and IM revied and edited the manuscript. All authors read and approved for publication.

## Data availability and statement

Source data for all the graphs included in this paper are available as supplementary data.

